# Dissociable Pupil and Oculomotor Markers of Attention Allocation and Distractor Suppression during listening

**DOI:** 10.1101/2025.05.19.654866

**Authors:** Xena Liu, Maria Chait

**Author notes:** **Corresponding Author:** Xena Liu, Ear Institute, University College London, 332 Gray’s Inn Road, London WC1X 8EE, UK; Maria Chait, Ear Institute, University College London, 332 Gray’s Inn Road, London WC1X 8EE, UK.

## Abstract

Emerging evidence suggests that ocular dynamics, pupil dilation, eye movements, and blinks, can serve as markers of task engagement. In this study, we examined whether these measures dissociate different perceptual states during an auditory attention task. Participants (Main and Control experiments; N = 34 and 27, both sexes) listened to tone sequences designated as ‘Target’, ‘Distractor’, and ‘Probe’, designed to isolate active attention from suppression of irrelevant input.

We focused on four physiological signals: pupil diameter (PD), pupil dilation rate (PDR), blink rate, and microsaccades (MS). Compared to a control condition, both Target and Probe intervals elicited sustained PD increases, blink suppression, and prolonged MS inhibition indicating extended attentional engagement. Interestingly, the Distractor interval also showed prolonged blink suppression and elevated PDR, suggesting active arousal to suppress irrelevant input. However, MS dynamics during Distractors were indistinguishable from control, implying that attention was not allocated to the distractors despite the heightened arousal. This dissociation supports the interpretation that MS is specifically modulated by attentional allocation, while PD and blinks also reflect general arousal or suppressive mechanisms.

Overall, these findings reveal that PD, MS, and blinking capture distinct aspects of cognitive control. Together, they offer a multidimensional framework for studying selective attention and distractor suppression in complex tasks.

**Significance statement:** Understanding an individual’s moment-to-moment state during demanding attention tasks is crucial for identifying perceptual challenges, explaining inter-individual variability, and diagnosing difficulties in special populations. Ocular measures, such as pupil size, blinks, and microsaccades, are gaining traction as objective, non-invasive indicators of perceptual and cognitive states. In this study, we demonstrate that, during a challenging auditory task requiring selective attention and active ignoring, different eye-related signals capture distinct aspects of listener state. Specifically, the dynamics observed during active attention differ systematically from those during active distractor suppression. These findings highlight the value of ocular metrics for assessment of cognitive performance and for disentangling the contributions of arousal and attention during complex tasks.

## Introduction

Successful listening in busy acoustic environments requires dynamic engagement of arousal and attention. It is increasingly understood that insights into these processes can be gleaned from eye-related measures such as the dynamics of pupil dilation, saccades, and blinking (e.g. Rolfs et al., 2008; Oh et al., 2012; Wang & Munoz, 2014; Wang et al., 2017; Maffei & Angrilli, 2018; Kobald et al., 2019; Zhao et al., 2024; Gehmacher et al., 2024; Cui & Herrmann, 2023; Herrmann & Ryan, 2024; Widmann et al., 2025; Herrmann et al., 2025).

Non-luminance-mediated pupil dilation (PD) is linked to activity in the locus coeruleus (LC), the primary source of norepinephrine in the CNS, and a key regulator of arousal (Wang & Munoz, 2015; Joshi et al., 2016). Baseline pupil size and stimulus-evoked phasic responses are thought to reflect tonic and phasic LC activity, respectively - where tonic activity indexes sustained alertness, and phasic activity reflects transient changes in arousal (Kahneman & Beatty, 1966; Beatty, 1982; Granholm & Steinhauer, 2004; Wang et al., 2014; Wang & Munoz, 2015; Wang et al., 2017; Milne et al., 2021).

Eye movements reliably decrease during periods of cognitive demand (e.g., Kobald et al., 2019; Walter & Bex, 2021; Herrmann & Ryan, 2024). Blinks also become infrequent and more strategically timed, often occurring during natural pause points (Orchard & Stern, 1991; Nakano et al., 2009; Hoppe et al., 2018; Chen et al., 2022). This pattern reflects attentional gating - suppression of saccades and blinks during cognitively demanding tasks helps conserve processing resources by minimizing the disruptive effects of visual transients. As such, ocular behavior serves as a sensitive indicator of moment-to-moment cognitive and perceptual states. Among these behaviors, microsaccades (MS) - small, involuntary fixational eye movements occurring at approximately 1–2 Hz (Rolfs, 2009) - are particularly relevant. Because MS are compatible with pupillometry, which requires sustained fixation, they offer a valuable metric in contexts that preclude free-viewing paradigms (Cui & Hermann, 2023).

The MS generation network - comprising the superior colliculus (SC), frontal eye fields (FEF), and visual cortex (Munoz & Istvan, 1998; Hafed et al., 2009, 2015; Rucci & Victor, 2015; Hsu et al., 2021) - is thought to underly automatic visual sampling. MS rates drop following the onset of salient stimuli, both visual and auditory (Hafed & Clark, 2002; Engbert & Kliegl, 2003; Wang et al., 2017), a phenomenon known as microsaccadic inhibition (MSI). MSI likely reflects rapid suppression of continuous sampling activity in response to attention-demanding input. Ongoing MS activity is further modulated by task load (Dalmaso et al., 2017; Lange et al., 2017; Xue et al., 2017; Badde et al., 2020; Kadosh et al., 2024). In the auditory domain, MS rates have been shown to decrease during periods requiring heightened attention (Widmann et al., 2014; Contadini-Wright et al., 2023; see also Abeles et al., 2020), suggesting that MS dynamics can serve as markers of instantaneous attentive engagement.

Here we contrast **active allocation of attention** with **active withdrawal of attention** to investigate their respective effects on ocular dynamics (Figure 1). Listening in complex auditory environments requires the focused allocation of attention to relevant auditory sources alongside deliberate suppression of distracting inputs (Tun et al., 2002; Nolden et al., 2018; Kattner & Ellermeier, 2020). Increasing evidence suggests that distractor suppression is an active, resource-intensive process, engaging neural systems distinct from those underlying top-down attention (Chait et al., 2010; Bidet-Caulet et al., 2010; Ahveninen et al., 2017; Wöstmann et al., 2022). This raises an important question: do ocular dynamics differ between active attention and active ignoring? Uncovering this relationship will offer deeper insight into oculomotor and pupillary markers of cognitive resource allocation, and clarify how arousal and attentional control interact under demanding conditions.

**Figure 1.**
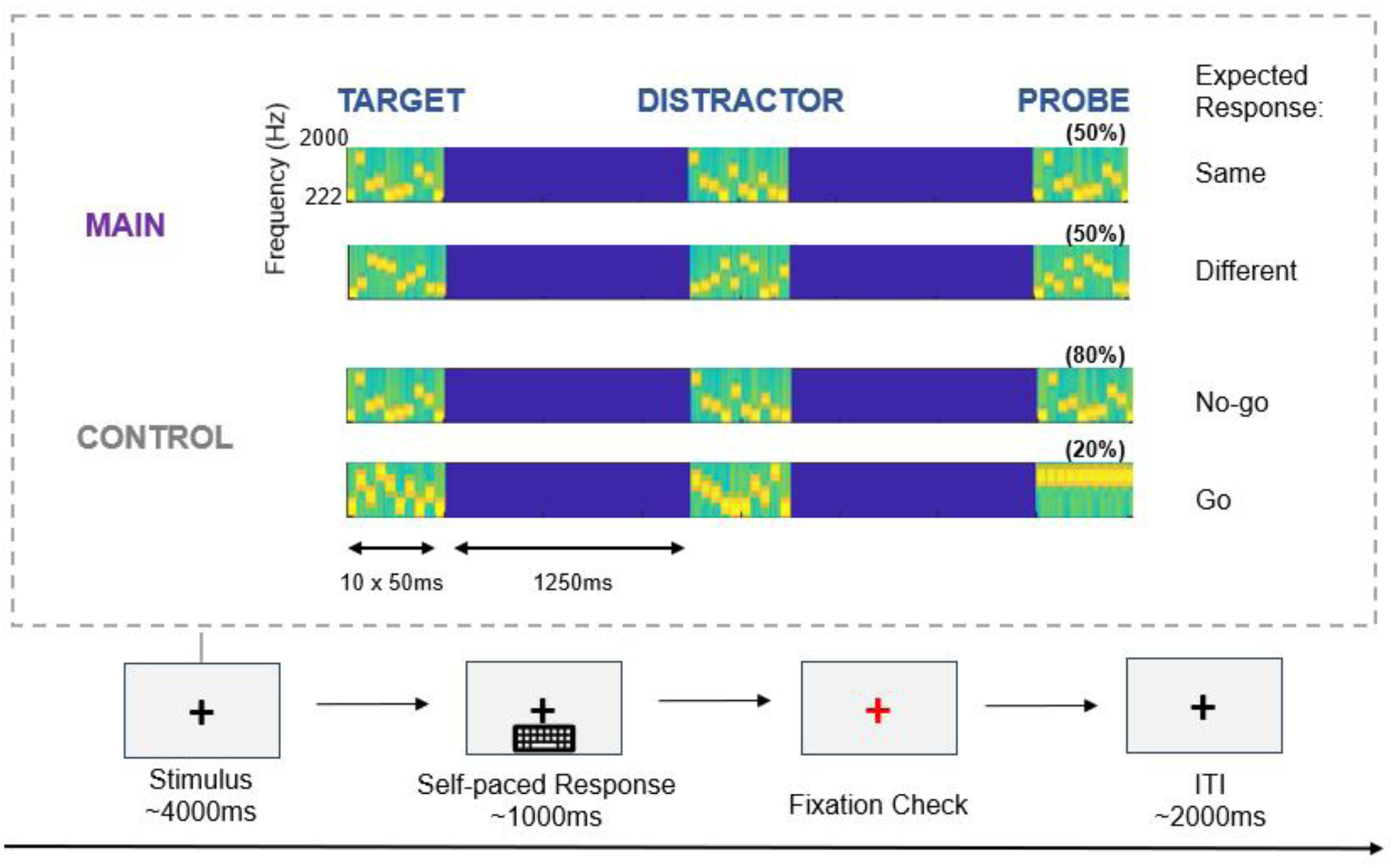
Stimulus structure (main and control experiment). The general trial structure (schematized in the bottom of the figure) consisted of a stimulus presentation phase, followed by a response phase (self-paced) where participants pressed one of two keyboard buttons to indicate their response, then a fixation check and an ITI of 2000ms. The stimulus period (lasting 4000ms) consisted of the successive presentation of a Target, Distractor and Probe sequence separated by silent intervals. In the Main experiment, participants determined whether the Probe was identical to the Target (50% of trials). In the Control Experiment, participants responded when the ‘Probe’ sequence consisted of a single repeating tone pip (20% of trials).

## Methods

### Participants

All participants were native or fluent English speakers, all reported normal hearing without history of otological or neurological disorders, and had normal or corrected-to-normal vision, with Sphere (SPH) prescriptions no greater than -3.50. All experiment procedures were approved by the UCL research ethics committee, and written informed consent was obtained from all participants. Participants were paid for their participation.

#### Main experiment

For the behavioral pilot: Ten participants aged between 18 and 46 were recruited. Data from 3 participants were excluded from analysis due to chance level performance. Therefore data from 7 participants are reported: mean age = 27.8, median = 26.0, SD = 7.69, 3 females.

For the eyetracking experiment, 35 participants aged between 18 and 60 were recruited. Data from one participant were excluded from the analysis due to excessive missing pupil data (> 50% blinks and/or gazes away from the fixation cross), resulting in 34 participants in the final dataset (mean age = 27.5, SD = 8.4, 24 females).

#### Control experiment

Thirty-one different participants aged between 18 and 60 were recruited. Data from 4 participants were excluded from analysis due to excessive missing pupil data (same criteria as the main experiment), resulting in a final set of 27 participants (mean age = 28.2, SD = 10.7, 19 females).

### Design and materials

To dissociate attending and ignoring, the stimuli (Figure 1) consisted of three consecutively presented tone-pip sequences. Participants were instructed to remember the *Target* sequence and later compare it to a *Probe* sequence. A *Distractor* sequence was presented in between, containing the same tones as the Target and Probe but in a different order. Thus, successful performance required listeners to actively suppress the intervening Distractor sequence. All experiment materials were generated and implemented in MATLAB and presented via Psychophysics Toolbox version 3 (PTB-3).

### Main experiment

Each trial (Figure 1) consisted of three consecutive tone sequences, separated by 1.25 s silent gaps, followed by a self-paced response period. Tone sequences were composed of 10 tones (each 50ms; 500ms total) selected from a pool of 20 tones, within the frequency range of 222 - 2000 Hz. Participants were instructed to remember the first tone sequence (‘Target’) and determine whether the third sequence (‘Probe’) is the same or different, whilst ignoring the middle sequence (‘Distractor’).

The tones comprising the Target were randomly selected from the frequency pool (without replacement) on each trial. The Distractor sequence was composed of the same 10 frequencies as the Target, but shuffled in order. The Probe sequence was either identical to the Target (No Change), or contained a switch in position of 5 randomly selected tones (but never the first or the last tone in the sequences to avoid primacy/ recency effects). That the ‘Distractor’ sequence was comprised of the same tones as the Target and Probe increased the difficulty of the task and encouraged participants to ignore the Distractor sequence to avoid interference in memory.

This design resulted in each trial’s stimulus presentation phase lasting for 4 seconds in total, followed by a self-paced keypress response period. Brief visual feedback on accuracy and reaction time was provided on the screen after each trial. Participants were incentivized with a potential bonus – they were informed that an extra payment of 1 pound per block would be awarded for all blocks that had more than 70% percent correct + fast responses (RT <1000ms; based on results of the pilot experiment described below).

All stimuli were generated offline and saved in 16-bit stereo wav format at a sampling rate of 44.1kHz. A different set of stimuli was generated for each participant. The main experiment session was composed of 4 blocks, each block containing 30 trials.

Before conducting the main experiment, a pilot experiment was run to assess behavioral performance (without concurrent eye tracking). The stimuli were identical to those described for the main experiment except 50% of the trials did not contain a Distractor sequence. These trials consisted of a Target, followed by a 3 sec silent gap and then a Probe. Trials with and without the distractor sequence were randomized in order and presented within the same block. The subjects’ task was identical to the main experiment. Participants completed 6 blocks (each 30 trials).

### Control experiment

In the control experiment, we aimed to measure ocular responses using the same trial structure as the main experiment, but under conditions that did not require focused attention, memory, or active ignoring. While we continue to refer to the sequences as "Target," "Distractor," and "Probe" for consistency and to indicate their positions within each trial, it is important to note that in the control experiment, neither the Target nor the Distractor held any behavioural relevance.

The triplet sequence structure was identical to that of the main experiment (Figure 1). The Target and Distractor sequences were generated in the same manner as previously described. The final sequence (Probe) was either a standard tone sequence (80% of trials) or a single, repeated tone lasting 500 ms (20% of trials). Participants were instructed to press a key whenever they detected the repeated tone. As such, attention was required only for the Probe sequence, and the task itself was perceptually simple and did not involve memory. The task was designed to maintain general engagement with the auditory stimuli without inducing significant cognitive load. As in the main experiment, participants completed four blocks of 30 trials each.

### Procedure

Participants sat in a dimly-lit soundproof testing room (IAC triple walled sound-attenuating booth), with their head fixated on a chinrest in front of a monitor (24-inch BENQ XL2420T with a resolution of 1920 × 1080 pixels and a refresh rate of 60 Hz). Ocular data were recorded from an infrared eye-tracking camera (Eyelink 1000 Desktop Mount, 1000Hz sampling rate; SR Research Ltd.) placed below the monitor at a horizontal distance of 62 cm. Auditory stimuli were presented binaurally through a Roland Tri-capture 24-bit 96 kHz soundcard connected to a pair of Sennheiser HD558 headphones, with volume levels adjusted to a comfortable level for each participant. The standard eye-tracking five-point calibration procedure was conducted before each experimental block. During the experiment, participants were instructed to fixate their eye gaze on a black cross in the middle of the screen, against a grey background. Gaze fixation was checked following each trial, and if fixation was detected correctly the next trial began after a 2-second inter-trial interval.

A practice block was run before the main task for all participants, all with the same practice stimulus set, which was different from the individual stimulus sets used in the experimental blocks.

### Data preprocessing and analysis

#### Pupillometry

Pupil data from the left eye were analyzed. During preprocessing, intervals with partial or full eye closures (e.g. during blinks) or where eye gaze was detected away from fixation (outside a radius of 100 pixels from the center of fixation) were discarded and treated as missing data. Participants with excessive missing data (>50%) were excluded from any further analysis. This applied to 1 participant in the main experiment and 3 participants in the control experiment.

Pupil data for each trial were epoched from -1 to 5s of trial onset. Trials with excessive missing data (>50%) were excluded from the analysis, and missing data in the remaining trials were recovered using linear interpolation. On average less than 2 trials per participant in each experimental block were excluded in this way. Outlier trials (trials with >50% data points 3 standard deviations away from the mean across trials) were also excluded. Less than 1 outlier trial per participant per block were excluded on average. Data were z-score normalized within each block and then epoched, time-domain averaged for each participant, and baseline corrected. Individual time series were then grand averaged to yield the data plotted in the Results section.

#### Pupil dilation event rate

Alongside the analysis of pupil diameter, we also analyzed the pupil dilation rate. This measure quantifies the incidence of pupil dilation events. Joshi et al. (2016) demonstrated a link between spiking activity in the LC, SC and inferior colliculus and pupil dilation, with spikes in all three areas correlated with subsequent pupil dilation events.

Following Joshi et al. (2016) and Zhao et al. (2024), events were defined as local minima with the constraint that continuous dilation is maintained for at least 0.1 s. The pupil events were extracted from the continuous data smoothed with a 0.15 s Hanning window. Similar to the analysis of MS events (described below), the detected pupil dilation events were then summed and normalized by number of trials and sampling rate to obtain an average at each timepoint, and then smoothed by a convolution window of α=1/50ms to obtain a time series of pupil dilation rate.

#### Blink extraction

Sharp drops in the raw eye movement data were marked as blinks, including full and partial (pupil size changes larger than 7 sd) eye closure movements. Identified blinks were stored and averaged at each timepoint to obtain a time series of blink rates (proportion of blinks per trial) for further analysis.

#### Microsaccades

Microsaccade (MS) detection followed the method proposed by Engbert and Kliegl (2003), which specified that MS events were extracted from continuous eye movement data based on the criteria: (1) a velocity threshold value of λ = 6 was used to multiply the median-based standard deviation of velocity distribution for each subject as the detection threshold; (2) above-threshold velocity lasting between 5 and 100ms; (3) events detected binocularly temporally overlap, with onset disparity <10ms; (4) the interval between successive microsaccades is longer than 50ms. For MS analysis, continuous eye movements for each trial were epoched from -2s to 5s relative to trial presentation onset.

For each participant, MS were detected according to the above criteria, based on the horizontal continuous eye movements. Event time series were then summed across trials and normalized by the number of trials and sampling rate. A causal smoothing kernel ω(τ)=α2×τ×e−ατ was then applied with a decay parameter of α=1/50ms (Dayan and Abbott, 2001; Rolfs et al., 2008; Widmann et al., 2014), paralleling a similar technique for computing neural firing rates from neuronal spike trains (Dayan and Abbott, 2001; see also Rolfs et al., 2008). The time axis was then shifted by the peak of the kernel window to account for the temporal delay caused by the smoothing kernel.

Following results from Rolfs et al. (2008) which demonstrated that MS amplitudes showed a decrease in close correlation to the time course of MS inhibition responses, we also extracted amplitudes of the identified MS events (distance between two positions on each MS trajectory) to assess whether MS amplitudes showed a similar response to attentional modulation. Distributions of MS amplitudes were constructed and compared between the main and control experiment.

#### Statistical analysis

For the behavioral data (in the pilot experiment), performance accuracy was computed for each condition in each subject (with and without distractor) and compared using a paired t-test.

In the main experiment, all eye data (pupillometry and MS) were first averaged across epochs to obtain a time series per subject. Correct and incorrect trials were also extracted based on the behavioral outcomes of each trial: for each subject, we selected the 30 correct trials with the fastest reaction times and 30 random incorrect (both “miss” and “false alarm” trials; on average, participants produced 45 incorrect trials overall, ranging from 25 to 68 trials). In cases where the number of incorrect trials was less than 30, the number of correct trials was directly matched to the subject’s total number of incorrect trials. The correct trials were selected by sorted reaction time to reduce the effect of chance on the selection of correct trials, as trials which participants were less certain of or were selected by guessing would likely have longer reaction times, compared to trials which they were more confident with their answers (see results). The pupillometry and MS data were then computed separately for the correct and incorrect trials and compared using a nonparametric bootstrapping test (1000 iterations with replacement; Efron and Tibshirani, 1994). At each timepoint differences between the two experiments were deemed significant if the proportion of bootstrap iterations that fell above or below zero was >95% (p<.05). A permutation analysis was performed to correct for multiple comparisons. For each comparison, all datasets from both conditions (main vs. control experiment) were pooled and randomly reassigned into two new groups, each containing an equal number of trials drawn from both original conditions. These permuted groups were then compared on a sample-by-sample (timepoint-by-timepoint) basis using the same statistical test as in the original analysis. For each timepoint, a binary outcome was recorded: 1 if the test yielded a significant difference (p < .05), and 0 otherwise. This permutation procedure was repeated 100 times. The proportion of permutations yielding significant results at each timepoint was used to estimate the False Discovery Rate (FDR). Timepoints in the actual data were considered to show reliable effects if the observed p-value was below .05 and the corresponding FDR estimate was below the threshold of .05.

## Results

### Control experiment

Figure 2A shows the performance in the control experiment (mean accuracy = 0.997, SD=0.005). The control task was specifically designed to require minimum computational/attentional resources and therefore high performance was expected.

**Figure 2.**
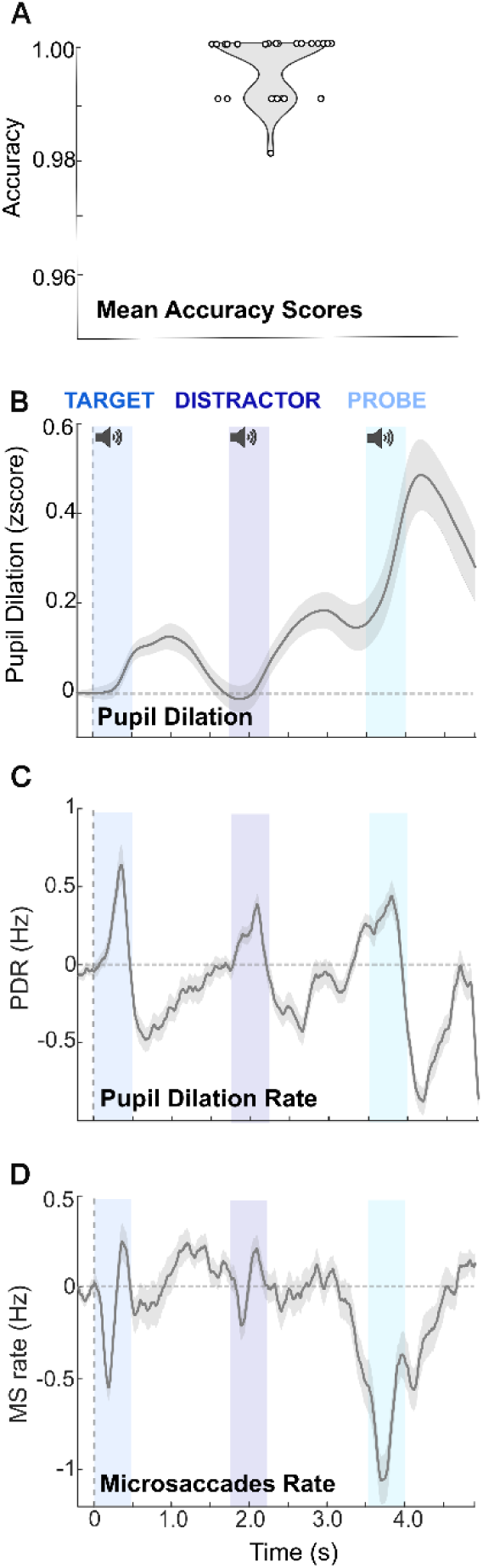
Control experiment results. **A**: Behavioral performance (accuracy) in the control task. Scattered dots represent data from individual participants. **B**: Mean pupil diameter (z-score) across participants, data are baselined according to [-0.5:0 s pre onset]. Blue shaded areas correspond to intervals of auditory stimulus presentation. **C:** Mean pupil dilation rate (Hz), data are baselined according to [-0.5:0 s pre onset]. **D:** mean MS rate, data baselined according to [- 0.5:0 s pre onset]. Error bars represent +/- 1STDE.

Figure 2B illustrates the task-evoked pupil dilation response in the control experiment. Pupil diameter exhibited peaks following the onset of each tone sequence, consistent with previous literature (Murphy et al., 2011; Liao et al., 2016; Wang et al., 2017; Zekveld et al., 2018), and likely reflecting sound-onset-evoked phasic arousal. The PD pattern following onset of the first two sequences (Target and Distractor) which did not require subjects to specifically attend to or ignore were relatively similar - gradually increasing to peak at approximately 1s after sequence onset. The PD amplitude reached its largest peak following onset of the Probe, which required most attentional engagement in this experiment. It is notable that this PD peaked approximately 750ms after the Probe sequence onset, which was earlier than in the first two sequences.

Analysis of PDR (Figure 2C) allows us to focus on the dynamics of phasic arousal. PDR peaked around 400ms following each tone sequence onset, with similar peak amplitudes following the three sequences.

Figure 2D illustrates the mean MS rate during the control experiment. MS rates first showed an abrupt microssacadic inhibition (MSI) response approximately 200ms after the Target sound onset, followed by a rebound. This prototypical dynamic, commonly observed in response to abrupt sensory events, has been hypothesized to reflect a rapid suppression of ongoing activity in the SC evoked by new resource-demanding sensory input (Engbert & Kliegl, 2003; Hafed & Clark, 2002; Wang et al., 2017). After this initial response, the MS rates rebounded to above baseline levels, and remained above baseline, potentially reflecting a release of attention as the sound was irrelevant to the task. A similar MSI and rapid rebound pattern was also shown following the Distractor but was overall smaller. After remaining around baseline level, the MS rates then started rapidly dropping prior to the onset of the Probe, reached its largest trough during presentation of the Probe, and gradually restored to baseline towards the end of the trial time course. Taken together, MS rates demonstrated a robust rapid MSI followed by rebound pattern reflecting immediate attentional draw and release in response to irrelevant sound stimulus onset, while MSI response to relevant and attentionally demanding stimulus was more enhanced and prolonged.

### Main experiment

Figure 3A displays the results of the behavioral pilot experiment where trials with and without a distractor were compared. The behavioral performance in the “no Distractor” trials - mean = 65.8, SD = 5.59 - was similar to what is expected based on previous work with this task (e.g. Bianco & Chait, 2023). A paired t-test confirmed that the average accuracy in the “no Distractor” condition was higher than in the “Distractor” condition (mean = 56.7, SD = 5.70) (t(12) = 3.01, p = .01), indicating that the presence of the Distractor sequence impaired performance, though due to the overall difficulty of the task the effect is small.

**Figure 3.**
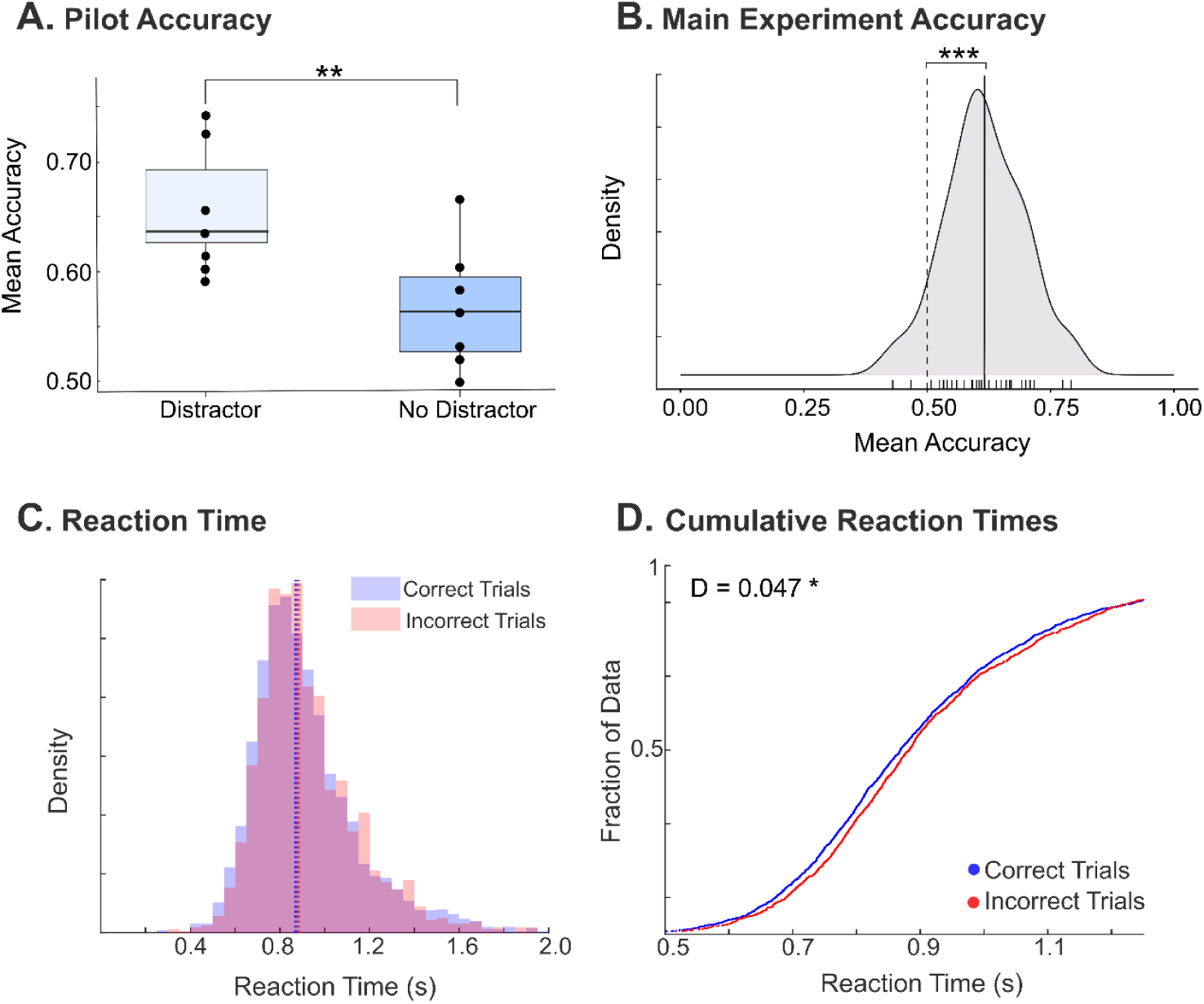
Main experiment - Behavioral performance. **A:** Auditory memory task accuracy scores in the pilot experiment for conditions in which a Distractor was presented, vs. not present. Filled circles represent individual subject data, and the horizontal lines within the boxplot represent group median. **B:** Distribution of accuracy scores of the behavioral task in the main experiment (during eye tracking). Individual data are shown on the x-axis. The solid vertical line represents group mean, and the dashed line represents chance level (accuracy of 0.5). **C:** Distribution of reaction times (RT) in correct and incorrect trials in the main experiment. **D:** Cumulative distribution of RT in correct vs. incorrect trials. As expected, RT in the Correct trials tended to be (slightly) faster than in incorrect trials, corresponding to a leftward shift in the cumulative distribution.

In the main experiment, we proceeded to include distractor sequences in all trials. Figure 3B. shows the overall distribution of accuracy outcomes. The mean accuracy across participants was approximately 0.62, which while relatively low, was significantly above chance (one sample t test: t(35) = 8.66, p<.001).

The overall mean reaction time (RT) across all trials and participants was 0.93 seconds (median = 0.92, SD = 0.15). Figure 3C demonstrates the distributions of reaction times for correct and incorrect trials in the main experiment respectively. Mean RT for correct trials was 0.928s (SD = 0.39), and 0.934s (SD = 0.26) for incorrect trials. Figure 3D shows the cumulative distribution of RTs in correct vs. incorrect trials. A Kolmogorov-Smirnov (KS) test showed that the distributions of RTs within correct and incorrect trials were different (D = 0.047, p =.026), consistent with a slight leftward shift (towards faster RT) for correct trials.

### Pupil size dynamics track the processing of tone sequences

Pupil responses in the main experiment are displayed in Figure 4A, shown against the data from the control experiment. Overall, PD in the main experiment exhibited larger deflections than those in the control experiment, specifically following the Target and preceding and following the Probe. This pattern aligns with the notion that heightened arousal was necessary at those time points – to process and encode the Target, and thereafter to prepare for the processing of the Probe. The latter being particularly demanding because it requires comparing the Probe to the memory trace of the Target, making a decision, etc. Interestingly, the PD data do not show any differences between the Control and Main experiment during or immediately following the Distractor sequence.

**Figure 4.**
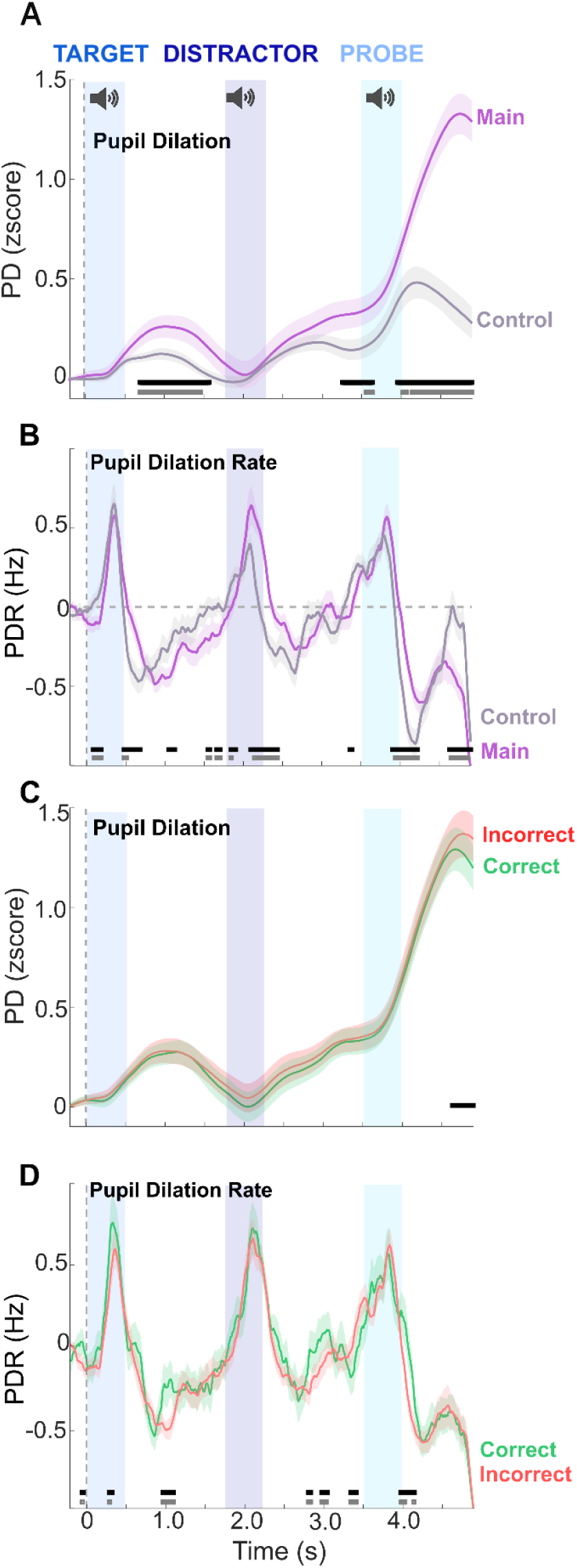
Pupil data in the main experiment. Purple lines in A and B correspond to results from the main experiment, and grey lines correspond to the control experiment results as shown previously. Green lines in C and D represent extracted correct trials in the main experiment and red lines represent incorrect trials data. Data are baselined according to [-0.5:0 s pre onset] time window. Error bars are +/- 1 STDE. Black horizontal bars represent timepoints where group differences are deemed significant (p<.05); grey horizontal bars below correspond to FDR corrected significance.

PDR, as a measure of instantaneous arousal, revealed differences following each of the sequences with slower decrease in PDR following the Target, Distractor and Probe in the main experiment relative to the control experiment (Figure 4B). This is consistent with participants maintaining instantaneous arousal longer following the three sequences in the main experiment. Notably this effect was most pronounced following the distractor sequence, suggesting increased instantaneous arousal during this period in the main experiment relative to the control.

Figures 4(C,D) plot PD and PDR data separately during correct vs. incorrect trials (see methods). There was no significant difference in PD dynamics between correct and incorrect trials except for a brief period at the end of the trial where correct trials show a PD decrease earlier than incorrect trials. This is consistent with the behavioral RT data (slightly faster in correct trials, see Figure 3D). For PDR, whilst noisy, the data indicate somewhat increased PDR across the whole trial duration during correct relative to incorrect trials, suggesting that (ultimately) correct responses are associated with increased instantaneous arousal.

### Demands on attention are associated with prolonged microsaccadic inhibition (MSI)

Figure 5A present MS dynamics in the Main experiment, against the data from the control experiment. The two conditions are characterized by a similar initial MSI response to Target onset. However, whilst the Control condition exhibits a subsequent rebound, in the Main experiment MS rate remains low for the sequence duration (500ms). Thereafter it returns to baseline, but remains lower than that in the control experiment, consistent with increased attentional load in the main task. A pattern similar to that observed during the Target is also then seen during the Probe sequence – a similar initial MSI is followed by sustained MS rate reduction in the main experiment, consistent with the enhanced attentional demands during that interval. Overall MS rates in the main experiment are lower than in the control.

**Figure 5:**
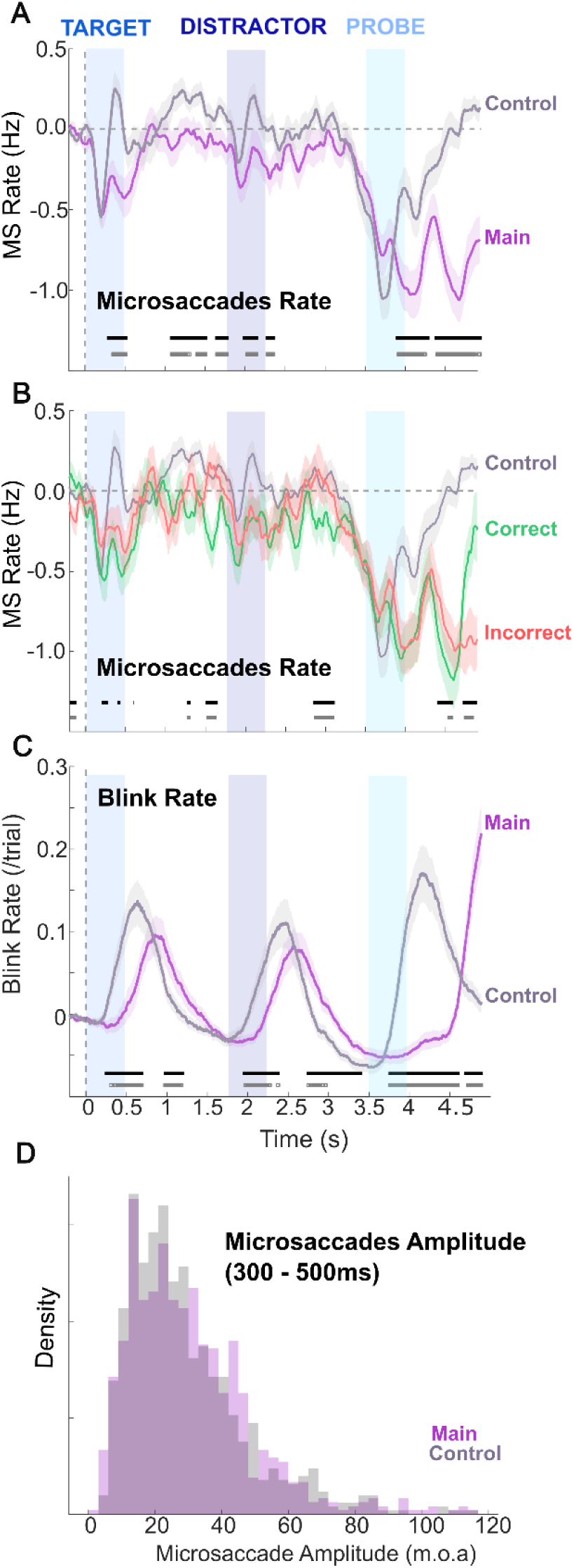
MS and blink data in the Main vs. Control experiments. **A:** MS rates. **B:** MS rate results after extracting correct(green) and incorrect(red) trials in main experiment. The significance bars below correspond to comparisons between correct and incorrect trials. Data are baseline-corrected according to [-0.5:0 s pre onset]. **C:** Blink rate, also baselined from [-0.5:0 s pre onset]. **D:** Distribution of amplitudes of MS events in Main vs. Control experiment extracted between 300 – 500ms of sound onset (i.e. during the presentation of the first sequence).

Figure 5B compares MS rates from correct and incorrect trials in the main experiment along with the MS rates (across all trials) in the control experiment. Though the significance pattern is noisy, MS rates were generally lower during correct trials compared to incorrect trials, consistent with the notion that heightened attention contributes to correct responses.

As an additional analysis, amplitudes of the MS events were extracted and compared for the two experiments (see methods). The plotted MS amplitudes in Figure 5D were extracted from 300 - 500ms of sound onset across trials. This time period was selected as a sample for this analysis as it corresponds to an interval where MSI dynamics in the main experiment differ substantially from those in the control. However following a KS test, no significant differences were found in the distributions of MS amplitudes between main and control experiment within this epoch (p = 0.13). This suggests that the experimental context of the main experiment did not elicit a distinct pattern of MS amplitude relative to the control condition.

### Blinks dynamics reflect attentional engagement

Blinks during the experiment were identified as events and extracted to produce a measure of blink rates throughout the trial time course. As illustrated in Figure 5C, in both the control, and main experiment blink rates increased shortly after the presentation of each sequence, consistent with previous reports (Fukuda, 2001; Oh et al., 2012; Bonneh et al., 2016; Kobald et al., 2019; Brych & Händel, 2020; Murali & Händel, 2021). These increases were earlier, and larger, in the control experiment in alignment with previous observations that blink incidence and timing are affected by the participants’ engagement with the task - blink rates only increased in the main experiment after the full duration of each sound sequence, especially for the Probe, where blinks were almost entirely suppressed until 500ms, after the end of the sequence presentation.

### Sequence-locked responses reveal distinct PDR/MS/Blink response patterns to the Distractor

Figure 6 focuses on sequence-locked responses, baseline corrected over 100ms prior to each sequence onset. As observed in Figure 6A top, increased PDR are seen in the main experiment following each of the 3 trial sequences. The most pronounced effect in terms of amplitude and extent is observed during the Distractor sequence. For the Target and Probe sequences, differences were primarily found during the post-sequence periods. As shown in the corresponding Difference plot, the differences during the distractor were largest, specifically during the presentation of the sequence (between 200-400ms). To assess potential latency differences between conditions, we performed a bootstrap-based analysis (see Methods), iterated 100 times, to estimate the distribution of divergence latencies across condition pairs (Main vs. Control) during the Target, Distractor, and Probe intervals. For each iteration, we calculated the latency at which the two conditions began to diverge significantly. The resulting distribution (Figure 6B) revealed that the earliest divergence occurred during the Distractor interval— approximately 200ms before significant differences were observed during the Target and Probe periods.

**Figure 6.**
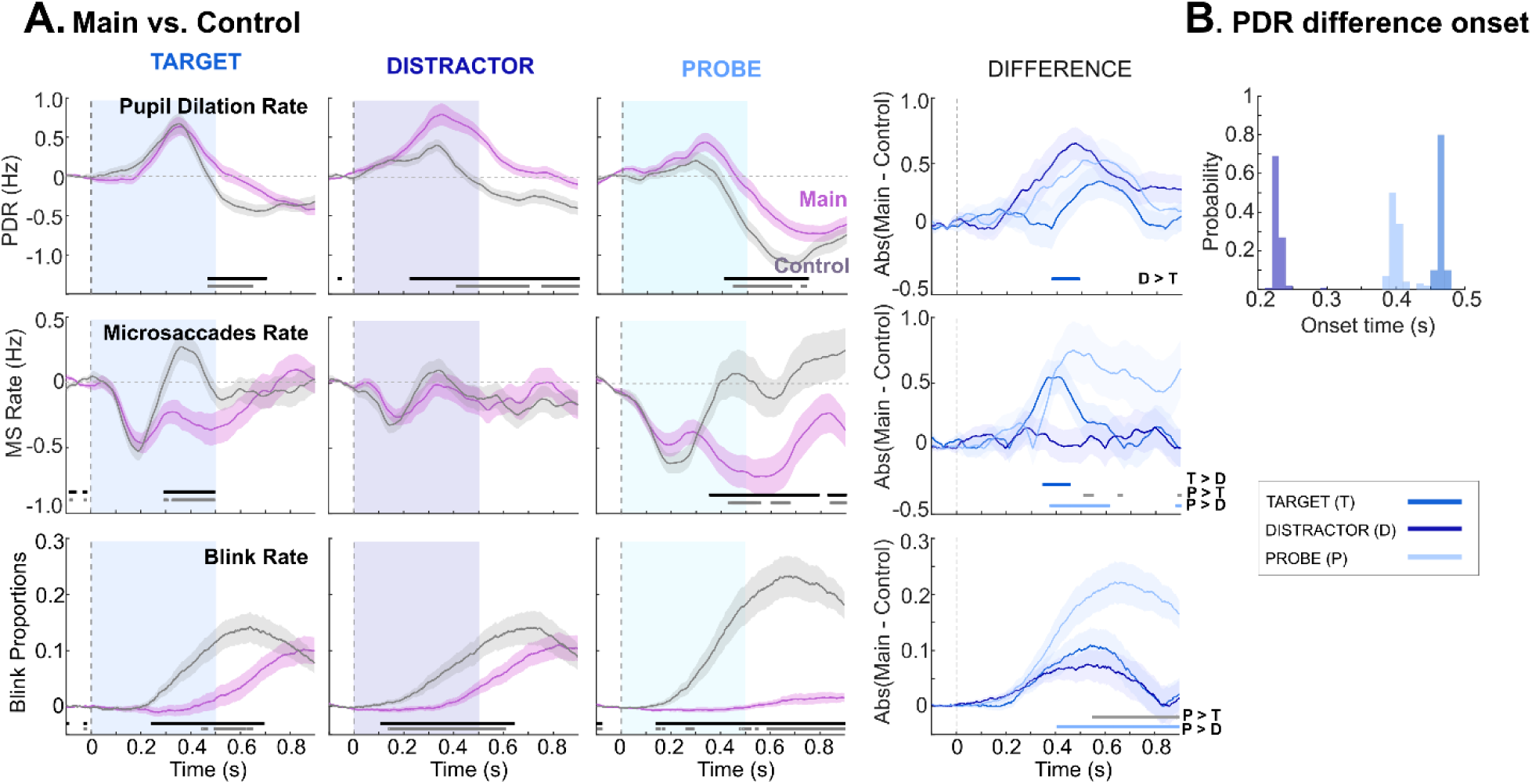
PDR, MS and blink rate data baselined [-0.1: 0 s] to the onset of each sequence. **A:** Comparisons between Main and Control experiments; Blue shading indicates time windows of sequence presentation (500ms). The right-most column (“DIFFERENCE”) illustrates group mean differences (absolute value) between main and control experiment. Horizontal bars indicate statistical significance. Black (p<0.05), Grey (FDR corrected); shades of blue (as indicated). **B:** A comparison of latencies of differences (between PDR data in Main and Control). Consistent with the plots in A, bootstrap resampling shows consistently earlier divergence for the Distractor relative to the Target and the Probe. Shaded areas around traces indicate +/- 1 STDE.

Figure 6A, middle, focuses on MS data. During the Target interval, the control experiment exhibited an MSI response with a trough at 200ms, followed by an immediate rebound, MS rate in the main experiment remained suppressed for the duration of the sequence (500ms), returning to baseline thereafter. A similar pattern was observed following the Probe, with a more extended suppression of MS consistent with the increased demands on attention during that period (encoding the Probe and retrieving memory of the Target). Notably, no differences between the control and Main experiment were observed during the Distractor sequence.

Figure 6A, bottom, focuses on blink rate data. In general, blinks were more suppressed in the main compared to the control experiment in all three time windows, especially during the stimulus presentation phases (0-500ms) where blink rates in the main experiment were close to baseline. Greatest differences in terms of both magnitude and latency were observed following the Probe sequence, where blinks in the main experiment were largely inhibited throughout the whole sequence epoch, consistent with highest time-locked demands on arousal.

Figure 7 illustrates sequence-locked PDR, MS, and blink rate comparisons between correct and incorrect trials in the main experiment. The data from the Control experiment (all trials) are also plotted for comparison. For PDR, differences between correct and incorrect trials were predominantly present during the Probe period, where (ultimately) correct trials were associated with increased instantaneous arousal. As for MS rates, significant differences between correct and incorrect trials were only observed during the Target sequence interval, where (ultimately) correct trials showed more enhanced MSI following sound onset. Blink dynamics did not differ between correct and incorrect trials across any of the analyzed intervals.

**Figure 7.**
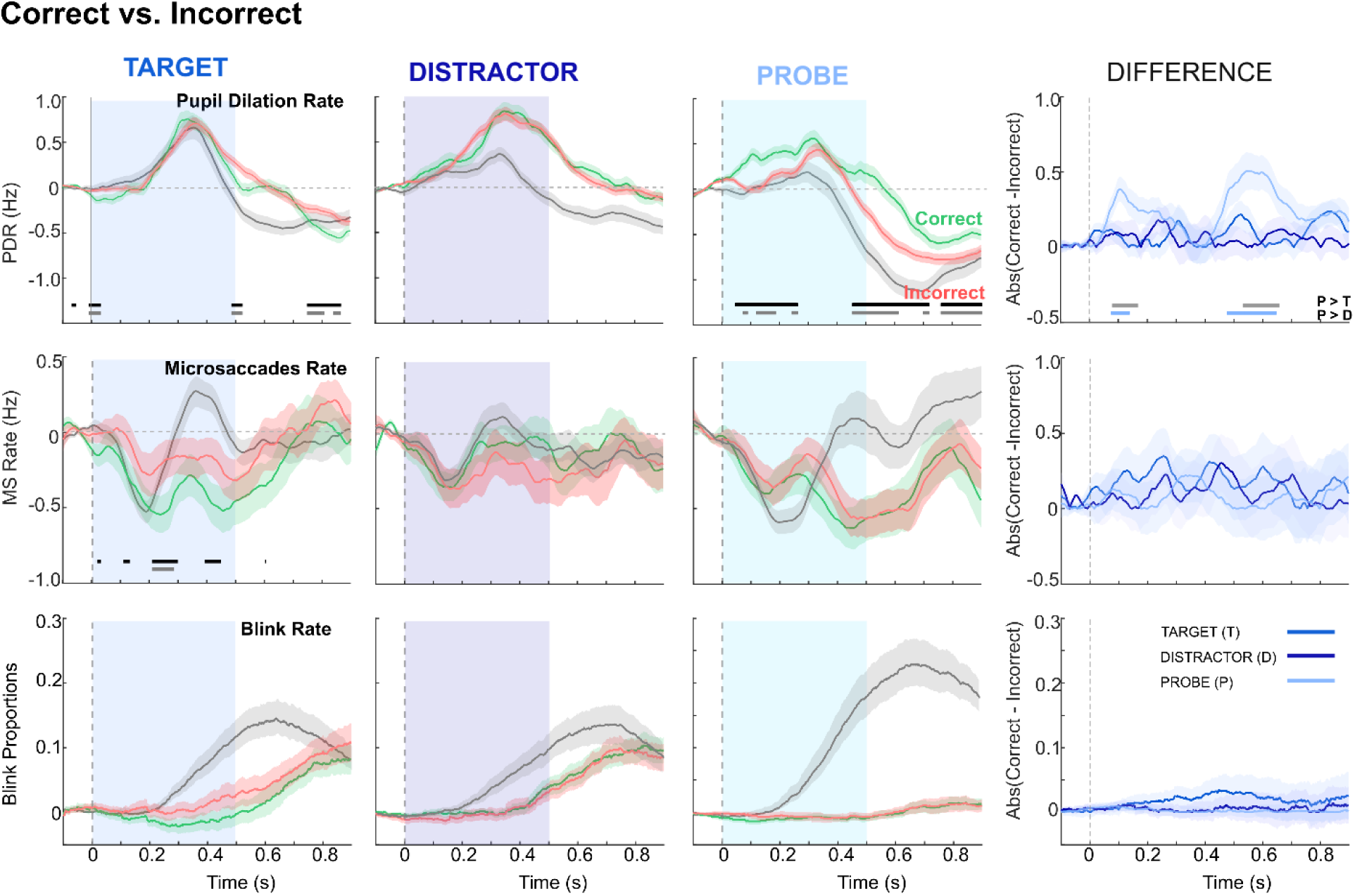
PDR, MS and blink rate data baselined [-0.1: 0s] to the onset of each sequence – comparison between correct and incorrect outcome trials from Main experiment. Data from the Control experiment (all trials; grey traces) are plotted for comparison. The right-most column (“DIFFERENCE”) illustrates group mean differences (absolute value) between correct and incorrect trials (green and red lines). Shaded areas around traces indicate +/- 1 STDE.

## Discussion

Ocular dynamics are increasingly recognized as valuable physiological proxies for task engagement, including in the auditory domain. Pupil diameter (PD) has consistently been linked to arousal and cognitive load (e.g. Beatty, 1982; Wang & Munoz, 2015; McGarrigle et al., 2017; van der Wel & van Steenbergen, 2018; Zekveld et al., 2018), while emerging evidence suggests that changes in eye movement dynamics, including saccades and blinks, reflect the instantaneous availability of attentional resources (Kobald et al., 2019; Contadini-Wright et al., 2023; Cui & Herrmann, 2023; Herrmann & Ryan, 2024; Herrmann et al., 2025). These findings align with the shared-resource framework, whereby the brain dynamically reallocates limited attentional and sensory resources across modalities in a push-pull fashion (Lavie, 2005): when task demands increase in the auditory domain, the visual system may be downregulated to reduce interference, including by suppressing saccadic eye movements and minimizing blinks, which introduce transient visual input that imposes additional processing demands. However, the neurocognitive mechanisms that underlie such cross-modal suppression remain obscure. Clarifying these processes is essential if ocular metrics are to be reliably used to assess auditory attention or to inform interventions.

We developed an auditory task designed to engage mechanisms of attentional allocation and distractor suppression. Our objective was to investigate how ocular indices reflect these cognitive demands during task performance. We focused on four physiological measures: pupil diameter (PD), pupil dilation rate (PDR) - as a temporally sensitive index of phasic arousal - blinks, and microsaccades (MS).

Our results show that PD, blinks, and MS display distinct patterns during distractor suppression.

### Top-down attention is associated with increased pupil dilation rate and prolonged suppression of microsaccades and blinks

Using data from a control experiment as a baseline, we demonstrate that attended sequences - specifically, the *Target* and *Probe* intervals - are associated with prolonged active processing. This is reflected in an enhanced and sustained phasic PD response, prolonged blink suppression, and an extended duration of MSI that persists throughout the entire sequence (500 ms). These observations are consistent with prior work in both visual and auditory domains. In the auditory domain, Widmann et al. (2014) reported prolonged MSI for target vs. non-target sounds, while Contadini-Wright et al. (2023) observed time-specific MS suppression during key word presentation in a speech-in-noise task.

In the visual modality, Kadosh et al. (2024) found that reduced MS rates during encoding, maintenance, and retrieval predicted better working memory performance. Similarly, we observed sustained MS rate modulation during the maintenance period, manifested as greater suppression in the main – relative to the control - experiment. Furthermore, although noisy, comparisons of correct vs. incorrect trials suggest that successful performance is linked to increased PDR and reduced MS rates.

However, a critical question remains: Is MS (and blink) suppression specific to attention, or do eye movements generally decrease in response to any situation that demands allocation of processing resources? The activity observed during the Distractor period is pivotal in addressing this question.

### Oculomotor Signatures of Distractor Suppression

Listeners were required to selectively attend to the Target and Probe sequences while ignoring the Distractor, which was constructed using the same tone-pips as the attended sequences. This design was adapted from a previous MEG study by Chait et al. (2010), which identified a sequence of early (<200ms post-onset) auditory cortical effects related to distractor suppression - comprising an attenuated onset response and the emergence of a subsequent component specific to distractor suppression. This component (see also Pomper & Chait, 2017) has been linked to selective attention and inhibitory control (Melara et al, 2002) and suggests that, when required, listeners possess a mechanism that allows them to selectively supress an interfering sound.

The component identified in response to auditory stimuli by Chait et al. (2010) and Melara et al. (2002) aligns with a larger body of work on the distractor positivity (Dp) response in vision (Gaspelin et al., 2023). The Dp is situated within broader theoretical frameworks suggesting that top-down attentional control operates through distinct facilitation and inhibition mechanisms (e.g., Bidet-Caulet et al., 2010; Gaspelin & Luck, 2018, 2019). While salient stimuli naturally capture attention, such capture can be prevented by proactively engaging suppression before attention is allocated. Converging evidence for these suppression mechanisms has emerged across multiple domains, including psychophysics (e.g., Gaspelin et al., 2015), eye-tracking (Adams et al., 2023), and brain imaging (Bidet-Caulet et al., 2010; Feldmann-Wüstefeld & Vogel, 2019; Gaspelin & Luck, 2018; Gaspelin et al., 2023), although this research has primarily focused on the visual modality.

A central question in the current study was whether ocular dynamics would show distinctive signatures of active distractor suppression. Specifically, given prior findings implicating MS rate reduction during periods of heightened attentional load, we asked whether MSI during the distractor period would be reduced, consistent with attention being successfully withheld, or potentially enhanced (i.e. resulting in a larger MS rate reduction), reflecting the possibility that MS-related circuits are not exclusively tied to attention per se, but also sensitive to resource availability due to the activation of top-down suppression mechanisms.

Using data from the control experiment, which required minimal attentional engagement, as a benchmark, we observed that blink suppression during the Distractor was prolonged, mirroring the pattern observed for the Target. Moreover, PDR was significantly increased during the Distractor period. This increase, quantified as the difference between the control and main experimental conditions, was in fact larger and earlier for the Distractor than for either the Target or the Probe. This pattern is consistent with a process of increased active arousal allocation, and suggests that the brain may engage in heightened suppressive mechanisms during the Distractor in order to protect task-relevant processing.

In contrast, MS dynamics during the Distractor period did not differ from those observed in the control condition; both exhibited a characteristic early MS inhibition (MSI) in response to sequence onset, followed by a rebound in MS rate. This pattern is consistent with attention being withdrawn from the Distractor and supports the view that MS dynamics are selectively modulated by attentional allocation, rather than by general arousal or non-specific cognitive effort.

Taken together, these findings suggests that the ocular metrics explored reflect different aspects of cognitive control, in line with the signal suppression proposal (e.g. Sawaki & Luck, 2010; Gaspelin & Luck, 2018). The increased PDR and prolonged blink suppression observed during the Distractor phase may reflect mechanisms of active suppression, potentially serving to prevent interference with working memory encoding. In contrast, the lack of change in MS dynamics supports the hypothesis that MS specifically index the allocation of attention, rather than general increases in processing demands.

### Oculomotor and Pupillary Markers of Cognitive Resource Allocation

Blinks, PDR and MS reflected distinct components of attention and suppression of distractors. Blink dynamics and PDR exhibited modulation across all three task phases with PDR showing most prominent modulation during the Distractor phase. In contrast, MS changes were observed exclusively during Target and Probe processing.

The neural circuitry underlying PD and MS is increasingly recognized as interconnected. The LC is closely linked with the FEF and SC, key regions involved in MS generation (Matsumoto et al., 2018). Micro-stimulation of the FEF and SC elicits pupil dilation and inhibit MS (Lehmann & Corneil, 2016; Joshi et al., 2016), consistent with these regions forming part of a broader network supporting attention and arousal regulation (Wang & Munoz, 2021; Sara & Bouret, 2012; Aston-Jones & Cohen, 2005). However, the precise functional roles and mechanisms of interaction among these structures remain under active investigation.

The observed dissociation between PDR and MS responses, particularly in the context of distraction, suggests at least partially functionally distinct underlying mechanisms (see also Schwetlick et al, 2025).

Perhaps most notable is the dissociation observed between blink and MS dynamics during the Distractor period. This divergence is unexpected, given that previous studies have reported similar inhibition patterns of spontaneous blinks and MS (Yablonski et al., 2017; Bonneh et al., 2016; Fried et al., 2014), consistent with the notion that blink suppression and MS rate reduction reflect similar processes: reducing visual processing so as to preserve a shared resource pool when demands on these resources are high.

Blinking and MS have been proposed to share inhibitory mechanisms via corollary discharges within the SC-FEF network but with at least partially distinct circuits: the SC is primarily involved in MS control (Sommer & Wurtz, 2002, 2006), while blink generation is more strongly associated with the facial and oculomotor nuclei and inferior regions of the FEF (Manning et al., 1983; Kato & Miyauchi, 2003; Coiner et al., 2019). The dissociation we observed is in line with this distinction and is consistent with a functional dissociation whereby blink suppression may be related to general arousal and computational demand, whereas MS suppression appears more specifically tied to attentional control.

Overall, these findings support the view that pupil dilation, MS, and blinking index distinct facets of cognitive resource allocation, offering a valuable framework for dissecting the neural correlates of attention and arousal in sensory processing, and for understanding the architecture of selective attention and cross-modal suppression.

## Conflicts of interest

The authors declared no conflicts of interest concerning the research, authorship, and/or publication of this article.

## Acknowledgments

This work was supported by RNID PhD studentship to XL. The funders had no role in study design, data collection, and analysis, decision to publish, or preparation of the manuscript.

## Data sharing

The data reported in this manuscript alongside related information will be available on OSF upon publication.

## Notes

### Competing Interest Statement

The authors have declared no competing interest.

